# censcyt: censored covariates in differential abundance analysis in cytometry

**DOI:** 10.1101/2020.11.09.374447

**Authors:** Reto Gerber, Mark D. Robinson

**Affiliations:** Department of Molecular Life Sciences, University of Zurich, Zurich, Switzerland; SIB Swiss Institute of Bioinformatics, Zurich, Switzerland

## Abstract

Innovations in single cell technologies have lead to a flurry of datasets and computational tools to process and interpret them, including analyses of cell composition changes and transition in cell states. The *diffcyt* workflow for differential discovery in cytometry data consist of several steps, including preprocessing, cell population identification and differential testing for an association with a binary or continuous covariate. However, the commonly measured quantity of survival time in clinical studies often results in a censored covariate where classical differential testing is inapplicable. To overcome this limitation, multiple methods to directly include censored covariates in differential abundance analysis were examined with the use of simulation studies and a case study. Results show high error control and decent sensitivity for a subset of the methods. The tested methods are implemented in the R package *censcyt* as an extension of *diffcyt* and are available at https://github.com/retogerber/censcyt. Methods for the direct inclusion of a censored variable as a predictor in GLMMs are a valid alternative to classical survival analysis methods, such as the Cox proportional hazard model, while allowing for more flexibility in the differential analysis.

## Background

Flow and mass cytometry are techniques to measure the presence of fluorochromes or isotopes conjugated to antibodies that are bound to specific cellular components at single cell resolution. Although cytometry can be considered an established method, recent developments enable the measurement of ever more markers simultaneously, resulting in a high-dimensional view for each cell [1, 2]. Although the number of measured features per cell is still much lower than in other single cell methods, such as single-cell RNA sequencing (scRNA-seq), the throughput is typically much higher with thousands of cells per second [1, 2]. An additional benefit of cytometry compared to scRNA-seq is the measurement at the protein level instead at the RNA level (since correlations between protein and mRNA expression can be low [3, 4]), although new cytometry-by-seq approaches (e.g. Cite-seq [5] and REAP-seq [6]) allow the simultaneous measurement of transcript and protein expression. The antibodies used in cytometry experiments are often chosen to discriminate several cell types by leveraging the biological knowledge about their protein expression (e.g. T-cells can be distinguished from other lymphocytes by the amount of CD3 they express). After obtaining the raw marker intensities per cell and preprocessing (including some or all of: Compensation, Quality assessment, Normalization, De-Barcoding, Filtering, Transformation [7, 8]), the first step is to discern cell populations. The historical approach for this clustering is manual gating, which requires an expert to choose thresholds of marker intensities to obtain (known) cell populations. Challenges around manual gating include lack of reproducibility and impracticability for high dimensionality [9, 10], which is why modern approaches try to overcome these limitations by either automatically choosing the best threshold to separate subpopulations (e.g. with *flowDensity* [11]), or by clustering cells using techniques such as *FlowSOM* (using a self organizing map) [12], *flowMeans* (k-means with cluster merging) [13] or *PhenoGraph* (based on a nearest-neighbor graph) [14]. Alternatives that do not strictly involve clustering include classifying cells based on an annotated reference dataset (e.g., linear discriminant analysis [15]).

After clustering or cell type assignment, the processed data contains a subpopulation label for each cell. The two classical analyses that can be performed are differential abundance (DA) and differential state analysis (DS) [16]. In DA, the (perhaps normalized) relative proportion of cells in a subpopulation per sample is tested for an association with additional information about the sample (e.g. control vs. treatment). The input data consists of a *cluster × sample* matrix of cell population abundances. In contrast, DS analyses organize the single cell data into *(cluster-marker) × sample* matrices, typically summarizing each subpopulation per sample with median marker expression; afterward, the summary is modeled against sample-wise annnotations for the association testing.

The R [17] package *diffcyt* [18] provides a framework for DA and DS for flow and mass cytometry. After preprocessing of the raw data, *FlowSOM* is (by default) used to (over)cluster cells into many small clusters representing potential rare cell populations [18]. DA can then be performed with well-known count-based methods *voom* [19], *edgeR* [20] or Generalized Linear Mixed Models (*GLMM*). Alternatives for differential discovery include, among others, *citrus* (overclustering, building of hierarchy, model selection and regularizations to get associations) [21], *cydar* (differential abundance on hypersphere counts, testing with Generalized Linear Models) [22], *CellCnn* (convolutional neural networks) [23] and *MASC* (Mixed-effects modeling of Associations of Single Cells) [24]. An important distinction is that, with *citrus* and *CellCnn* on one side and *diffcyt* and *MASC* (and *cydar*) on the other, the association testing is “reversed”: for *diffcyt*, the cell population (relative) abundances are represented in the statistical model as the response, whereas in *citrus* and *CellCnn*, the abundances are treated as a covariate. The reversed approach allows for more flexibility in the experimental setup since it allows to include additional covariates, such as batch or age, to be directly adjusted for [16], and *diffcyt* was shown to compare favourably in terms of sensitivity and specificity across several test cases [18].

Cytometry samples from clinical studies often contain additional patient data, such as treatment group (e.g., control vs. treated), age or survival time. DA with a binary variable (e.g. control vs. treated) can be seen as the “classical” case in cytometry. Of particular interest is whether a cell subpopulation is more abundant in one experimental condition compared to the other, which could be indicative of the effectiveness of a treatment. If an association with a continuous variable (e.g. age) is of interest, the modeling and testing are similar to the binary case and often the same methods can be used, since linear models underpin the statistical framework. If a time-to-event variable (e.g. time to an event, such as death or recurrence of disease) is considered, there is a need to use different methods altogether. The problem with time-to-event variables is a purely practical one caused by events that are “censored”, i.e., they are not fully observed but only a minimum (or maximum) is known.

An example of cytometry data of a clinical study can be found in the FlowCAP IV (Flow Cytometry: Critical Assessment of Population Identification Methods) challenge [25]. 13 marker intensities of PBMC samples of 383 patients linked to time to progression to AIDS from HIV+ were measured with flow cytometry, with the objective to find cellular correlates that predict survival [25]. At the time, the two best performing methods, (*FloReMi* [26] and *flowDensity/flowType/RchyOptimyx*), both relied on classical survival analysis methods in the association testing step, such as the Cox proportional hazard model [27], where the censored variable is modeled as the response and the subpopulation abundance as the predictor.

Meanwhile, the performant frameworks for cytometry analysis that have been shown to perform well with completely observed data (e.g. *diffcyt* [18]) cannot directly handle censored data; in particular, a censored predictor should not be treated as fully observed, since it can lead to a bias [28]. Removing incomplete samples can be a workaround, but is inefficient for high censoring rates and might lead to a bias as well [29]. Thus, the goal of this work is to investigate how to best directly include a *censored predictor* in the modeling framework, which itself is an under-researched area compared to survival response models. The following are noteworthy: Rigobon *et al.* described basic issues that arise from censored covariates [28]; Tsimikas *et al.* developed a method based on estimating functions for generalized linear models [30]; Taylor *et al.* described two methods based on multiple imputation [31]; Qian *et al.* developed a threshold regression approach [32]; Atem *et al.* developed methods based on multiple imputation in a bootstrapping setup [33].

In the following, we describe an extension to the linear model approach to DA in *diffcyt* that allows to directly include random right censored time-to-event variables as a covariate using methods based on multiple imputation. More specifically, risk set imputation and Kaplan-Meier imputation (imputation based on the Kaplan-Meier estimator of the survival function) from Taylor *et al.* [31] and the conditional multiple imputation (imputation based on the mean residual life) from Atem *et al.* [33] are included. A simulation framework was developed to evaluate basic properties of the model as well as differential discovery performance in the context of cytometry. The dataset from the FlowCAP IV challenge was re-analysed according to the *diffcyt* workflow with the censoring-specific methods to highlight real world applicability.

## Results

In order to test the performance of the included methods that handle censored covariates, two simulation studies were performed, the first exploring basic properties in a simplistic model and the second embedded in the situation of differential discovery performance when considering a single cell dataset with multiple subpopulations.

### Basic simulations

In the basic simulation, counts (*Y_j_*) for a sample *j* were modeled as binomially distributed with a GLMM association with two covariates, one censored (*T_j_*) and the other binary (*Z_j_*), via a logit link function with regression coefficients *β*:

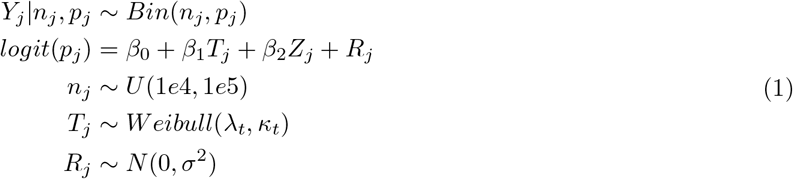

where *R_j_* represents an observation-level random effect to model overdispersion and *n_j_* is the total number of cells in a sample. For further details, see the Methods section.

Results of the basic simulations are shown in Figure 1 for three different censoring rates (30%, 50%, 70%) for a sample size of 100 with 100 repetitions per condition. Four different evaluation criteria are considered: raw bias 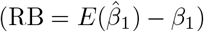, coverage rate (CR, proportion of confideJnce intervals that contain the true value), confidence interval (CI) width and root mean squared error 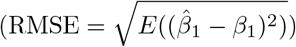. For a multiple imputation method to be considered “randomization-valid”, it should have no bias and a CR close to the specified proportion (in this case, 0.95) [34]. If a method is randomization-valid, the average width of the CI is another important criterion that represents statistical efficiency. On the other hand, the RMSE is an indicator of the precision of the estimation as it combines the variance and the bias 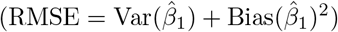. For increasing censoring rates, the RB (top row in Figure 1) for methods Kaplan-Meier imputation (*km*), Kaplan-Meier imputation with an exponential survival function tail (*kme*), risk set imputation (*rs*) and predictive mean matching (*pmm*) increases slightly, while for the other methods, it remains constant although the RMSE increases. In particular, the RB for those four methods is positive under all conditions, indicating overestimation. This observed bias is quite consistent across different simulation conditions (See Supplementary Figures S1-S4) although only for a low regression coefficient of the censored covariate does it become pronounced (Supplementary Figure S2). The CR (second row in Figure 1) is for all methods close to the expected value of 0.95 and taken together with the RB (in general close to zero) confirms the randomization-validness of the methods under most of the tested simulation conditions. The CI width (third row in Figure 1) for *km*, *kme* and *rs* has a nearly equal spread across all conditions while for the remaining methods, it increases with increasing censoring rate. Since the RMSE (bottom row in Figure 1) is a combination of the variance and the bias of an estimate it summarizes row 1 and 3 of Figure 1. So even though the estimates from *km*, *kme* and *rs* are slightly biased, their RMSE is lower compared to the other methods since their variance is lower.

**Figure 1.**
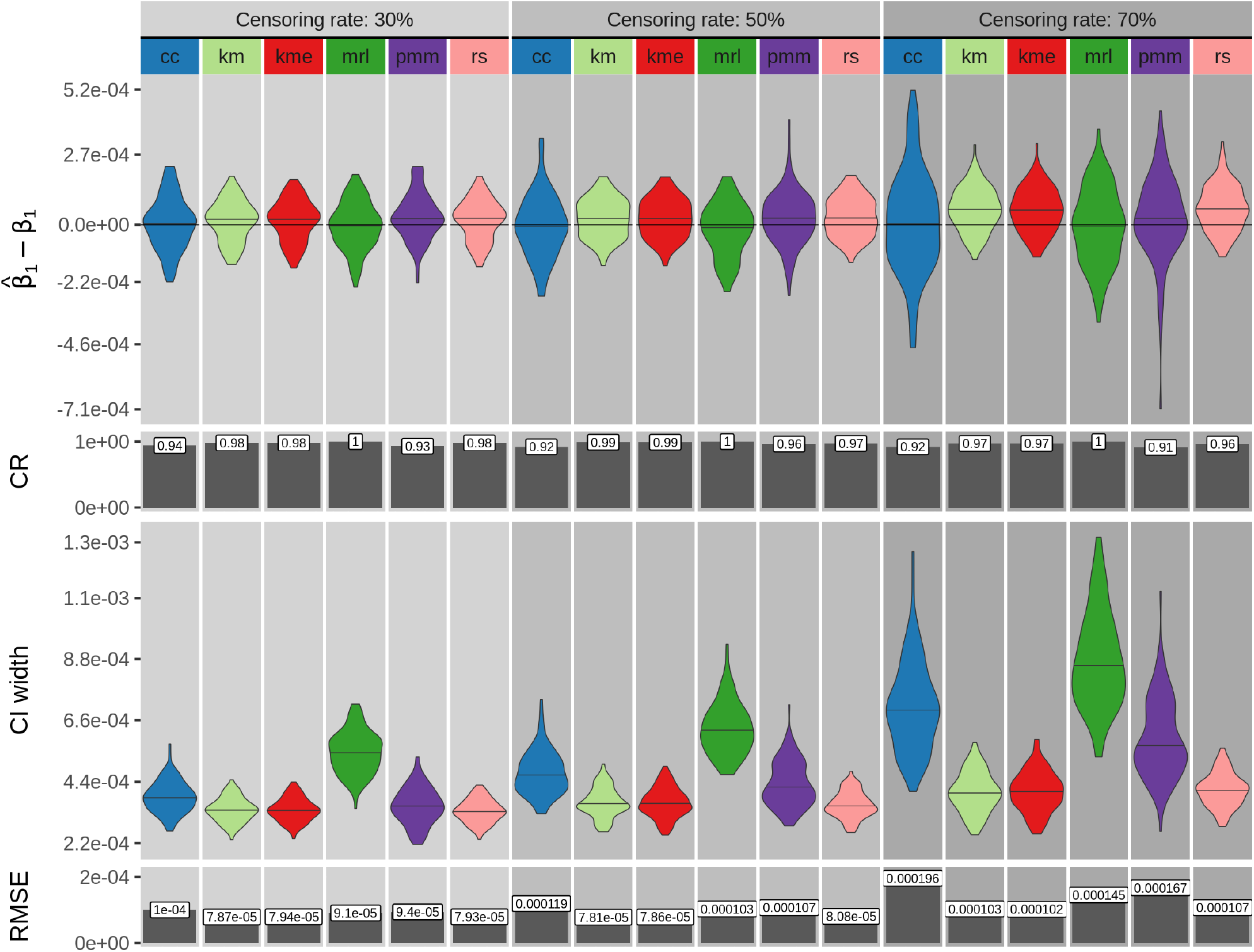
Single cluster simulation results for a sample size of 100 for censoring rates of 30%, 50% and 70%. Shown are four measures calculated from 100 simulation repetitions: difference of the estimated regression coefficient 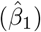 and its true value (*β*_1_), coverage rate (CR), confidence interval (CI) width and root mean squared error (RMSE). *cc*: complete case analysis, *km*: Kaplan-Meier imputation, *kme*: Kaplan-Meier imputation with an exponential tail, *mrl*: mean residual life imputation (conditional multiple imputation), *pmm*: predictive mean matching (treating censored values as missing), *rs*: risk set imputation. Other parameter values are: true regression coefficient *β*_1_ = −1*e* − 4, number of multiple imputations = 50 and the variance of the random effect = 1.

The distribution of p-values under the null simulation is for a low censoring rate uniform for all methods except *mrl* whose distribution is shifted towards 1 (Figure 2). For increasing sample sizes, the p-value distributions of all methods (except *cc*) shift towards 1, suggesting they become more conservative. The distribution for *cc* on the other hand shifts slightly towards 0.

**Figure 2.**
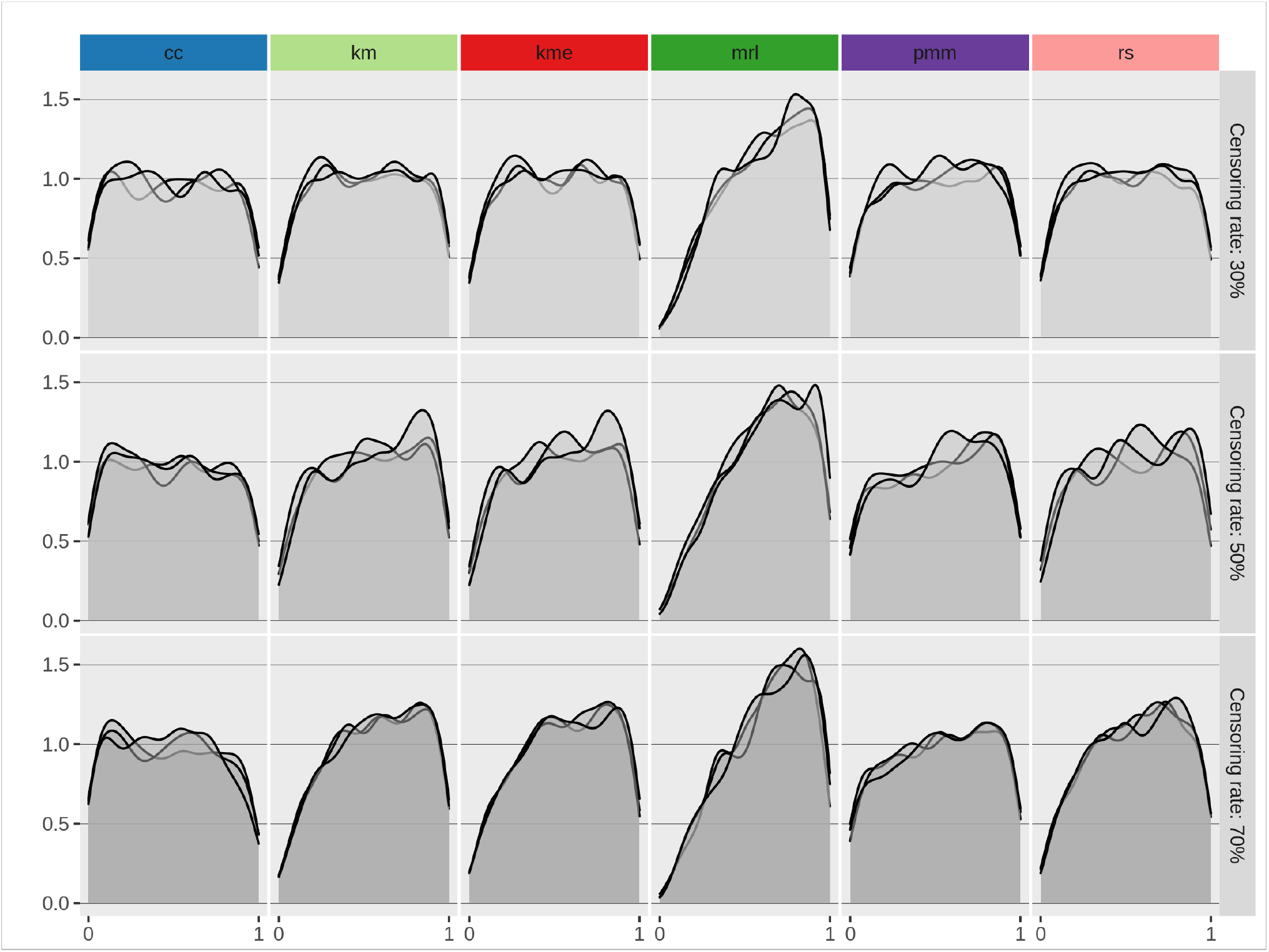
Single cluster simulation p-value distribution under the null model for three different censoring rates (30%, 50%, 70%). *cc*: complete case analysis, *km* Kaplan-Meier imputation, *kme* Kaplan-Meier imputation with an exponential tail, *mrl* mean residual life imputation (conditional multiple imputation), *pmm* predictive mean matching (treating censored values as missing), *rs* Risk set imputation. Each line represents 1000 repetitions.

Taken together, these results show that no tested methods stand out as being uniformly underperforming, but none is remarkably outperforming compared to its competing methods.

### Simulations modeled from real data

Figure 3 depicts a schematic of the simulation procedure for the multiple cell population scenario. Based on a real dataset clustered into cell populations (e.g. data from FlowCAP IV clustered with *FlowSOM*; Figure 3a), a Dirichlet-multinomial (DM) distribution is fit to the *cluster × sample* matrix of abundances (Figure 3b). To insert a known association, the obtained concentration parameters 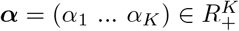 are then adjusted to include an association with a continuous (and later, censored) and a binary variable (Figure 3c, Eqn.3). The sizes (second parameter of DM) are kept the same. For subpopulation *i* ∈ {1*…K*} and sample *j* ∈ {1*…N*} the counts of a sample *Y_j_* ∈ *N^K^* with size *n_j_* ∈ *N* are distributed according to

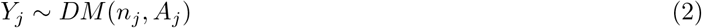

with the matrix of concentration parameters 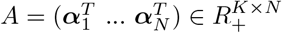 dependent among others on the continuous covariate *T_j_* and the binary covariate *Z_j_*:

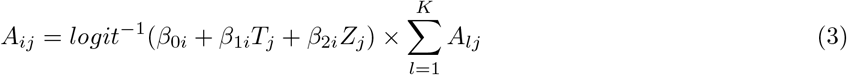

where the *β* parameters are the regression coefficients. A new dataset is then simulated with the adjusted parameters(Figure 3d). For further details, see the Methods section.

**Figure 3.**
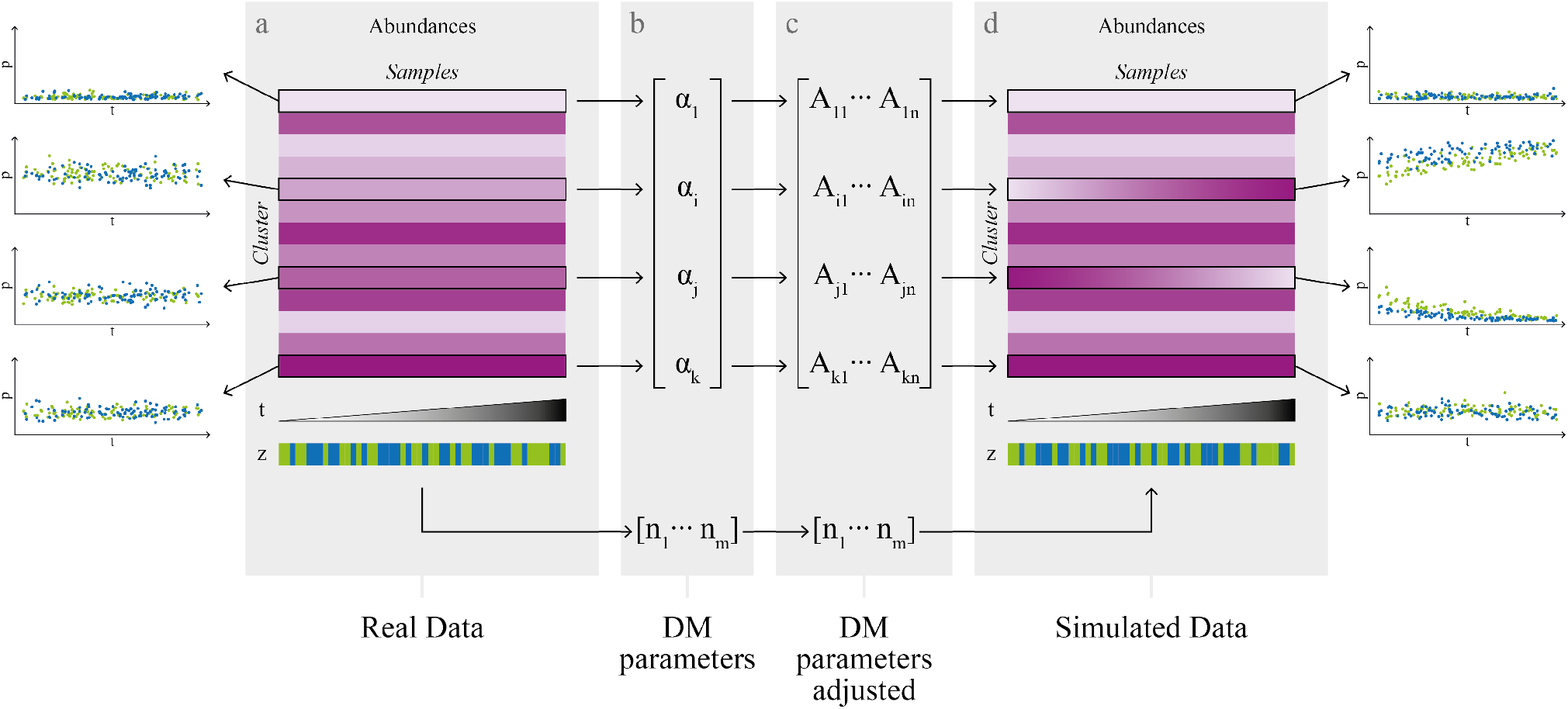
Simulation schema for multiple cell populations. (a) Starting with a *cluster × sample* matrix of abundances from a real dataset (b) a dirichlet-multinomial (DM) distribution is fitted. (c) The DM parameters are expanded and adapted to include an association of the abundances with a continuous covariate *t* and a binary covariate *z*. (d) A new dataset is simulated from the new parameters.

When two covariates are present, one option is to test for an association of the cell population abundance with the censored covariate (i.e. by testing if the regression coefficient of the censored covariate *β*_1_ = 0 in Eqn. 3) while also accounting for the binary covariate. In Figure 4 the TPR-FDR (true positive rate versus achieved false discovery rate) curves for the detection of true association between cell population abundance and survival time are shown for three different censoring rates and four different sample sizes.

**Figure 4.**
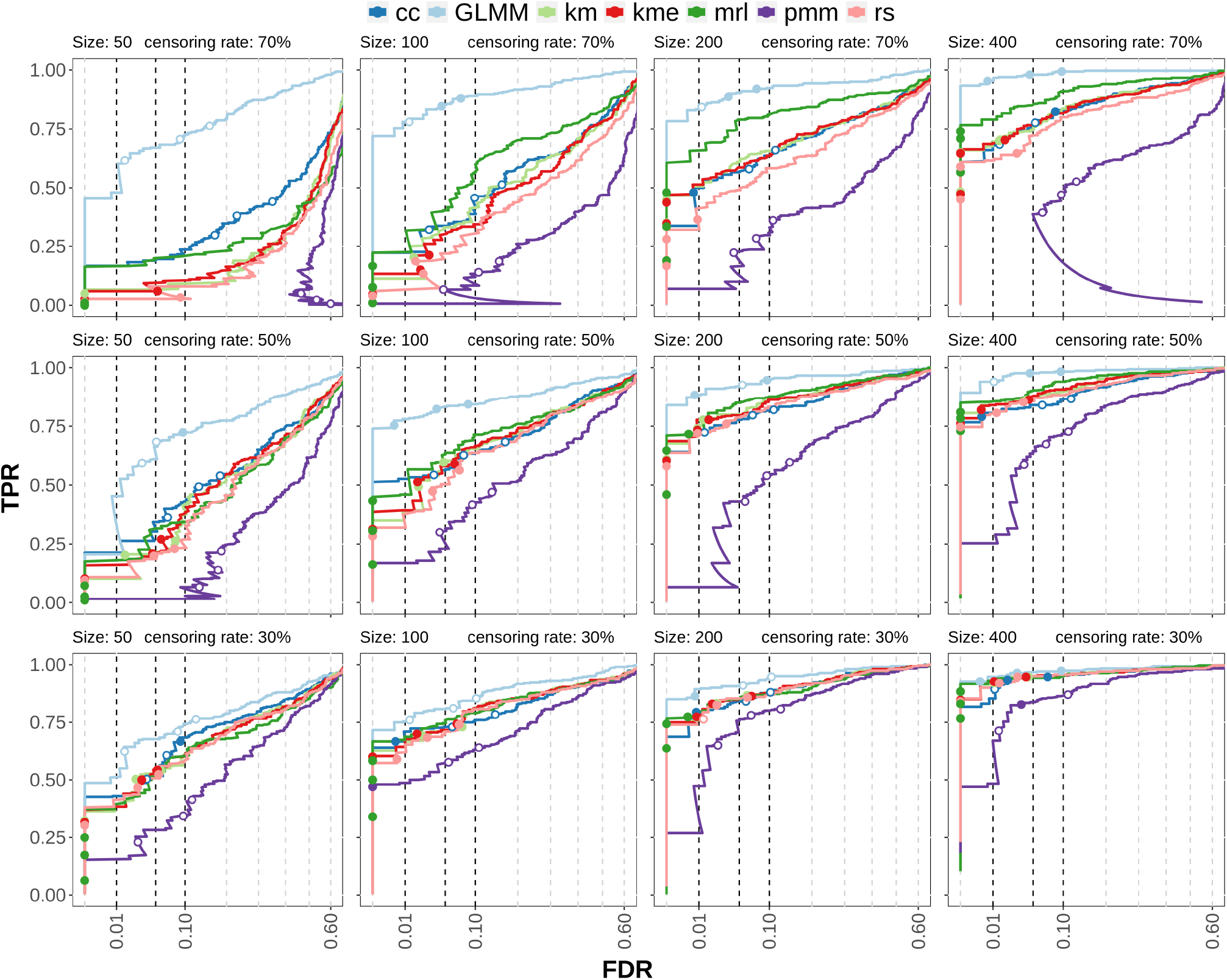
Multiple cluster simulation results testing for the association of the censored covariate. TPR-FDR curves for censoring rates of 30%, 50% and 70% (rows) and samples sizes of 50,100,200 (columns). The x-axis is square root transformed. *cc*: complete case analysis, *km*: Kaplan-Meier imputation, *kme*: Kaplan-Meier imputation with an exponential tail, *mrl*: mean residual life imputation (conditional multiple imputation), *pmm*: predictive mean matching (treating censored values as missing), *rs*: Risk set imputation. *GLMM* uses the (unobserved) ground truth of the survival time and can be considered to be the maximum possible performance of the other methods.

The method *GLMM* is the generalized linear mixed model method from *diffcyt* using the true (but unobserved) survival times and is included as a control, since it represents the maximum performance that could be achieved if the data were fully observed. It is not dependent on the censoring rate, so it can also be seen as a qualitative comparison of the simulation variability for a given sample size. *pmm*, on the other hand, can be considered to be a quasi-negative control, since it treats censored values as missing (leading to increased uncertainty about the data); thus, it highlights the gain in information from including censored values versus treating them as missing. In contrast, *cc* only keeps the “best” samples (the ones that are observed), which leads to more certainty about the data (at the cost of less data and potentially biased estimates).

Not surprisingly, lower sample sizes and increased censoring rates result in lower sensitivity. For a censoring rate of 30%, the differences in performance between the methods are minimal, independent of the sample size. For high censoring rates (70%), the differences between the methods are more prominent but decrease again for large sample sizes (400). *pmm* has overall the lowest sensitivity and poor error control; this is especially pronounced at high censoring rates leading to TPR-FDR curves with high FDR at low TPR. On the other hand, *cc* shows moderate sensitivity but the error control is poor for both high censoring rates and small sample sizes. *rs*, *km*, *kme* have in general a moderate sensitivity and good error control while *mrl* has good sensitivity and decent error control. Especially for high censoring rates, *mrl* outperforms other methods in terms of TPR.

To summarize: The censoring-specific methods have in general good error control, but especially for high censoring rates, result in lower sensitivity at a given p-value threshold (e.g. 0.05) than cc (which has poor error control).

The second option is to test for the association between the binary covariate and the cell population abundance (i.e. by testing if the regression coefficient of the binary covariate *β*_2_ = 0 in Eqn. 3), in the presence of a censored covariate. The TPR-FDR curves in this scenario (Figure 5) show clear differences compared to the testing for the association with the censored variable. *GLMM* is again the unrealistic control while *ncGLMM* is based on *GLMM*, but excludes the censored covariate in differential testing. It could therefore be seen as the ad-hoc solution when a censored covariate is present but not of interest and one decides to neglect the possible effect of the second covariate on the response. Two main differences compared to the association testing of the censored covariate is that *cc* and *mrl* have low sensitivity, even lower than *pmm* in many cases. The best performing methods are *km*, *kme* and *rs*, which often have similar sensitivity and error control. In many cases, they have a higher sensitivity than *ncGLMM* indicating that there is a benefit of accounting for the censored covariate instead of discarding it. Comparing the error control between Figure 4 and 5 shows that in the binary covariate association testing, the error control of the censoring-specific methods is often closer to its expected values than in the censored covariate association testing.

**Figure 5.**
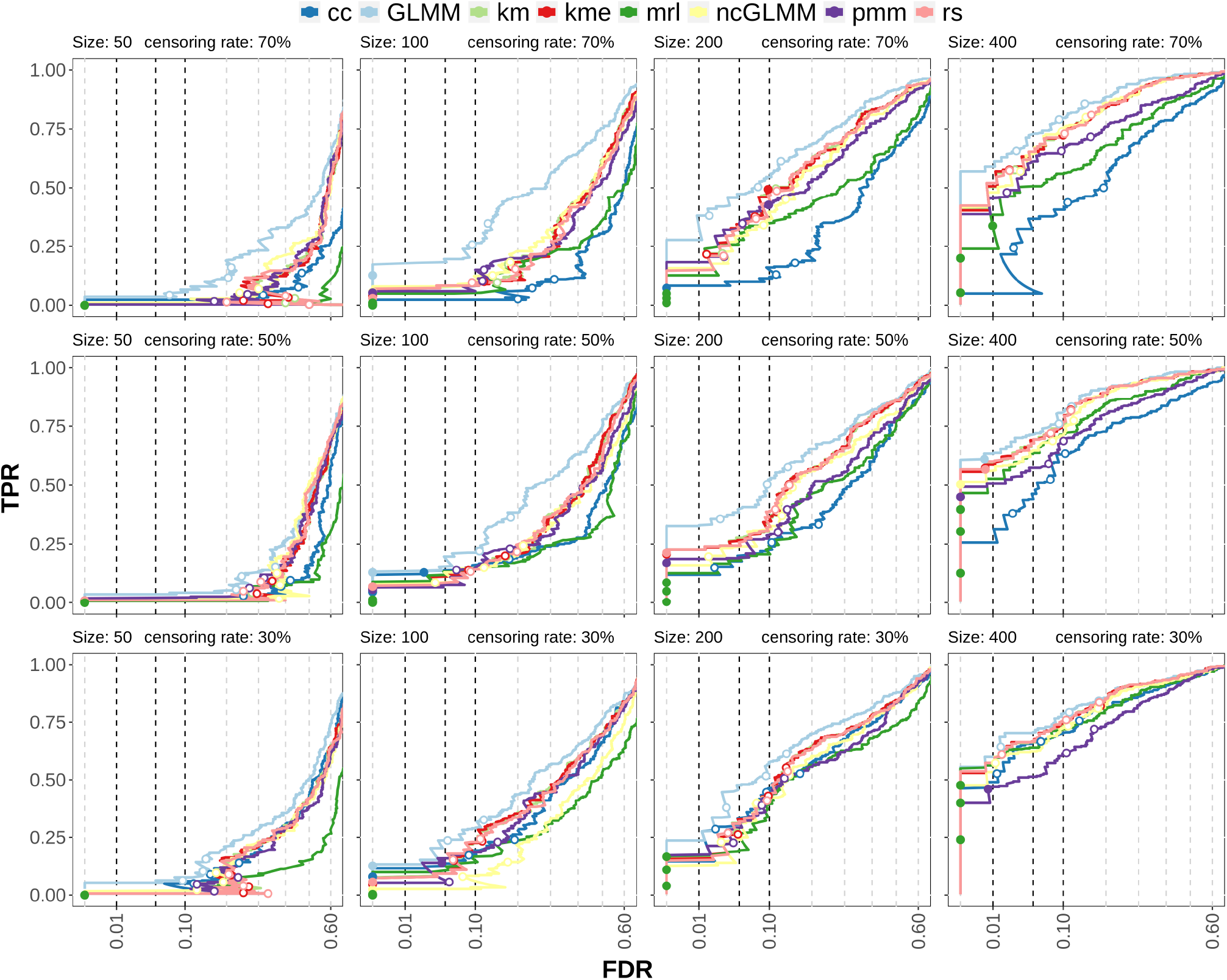
TPR-FDR curves of multiple cluster simulations with testing for the association of the binary covariate. Rows are censoring rates (30%, 50% and 70%) and columns are samples sizes (50, 100, 200 and 400). The x-axis is square root transformed. *cc*: complete case analysis, *km*: Kaplan-Meier imputation, *kme*: Kaplan-Meier imputation with an exponential tail of the survival function, *mrl*: mean residual life imputation (conditional multiple imputation), *pmm*: predictive mean matching (treating censored values as missing), *rs*: risk set imputation. *GLMM* uses the (unobserved) ground truth of the survival time and can be considered to be the maximum possible performance of the other methods. *ncGLMM*: same as *GLMM* but uses only the binary covariate for fitting and testing.

An alternative simulation scenario with only one censored covariate was modeled as well to compare censored-covariate methods with the Cox proportional hazard model [27] (by maintaining the simulated associations, but switching the response and the covariate in the statistical model). The results indicate similar performance in terms of specificity and error control for the Cox proportional hazards model and the censored covariate regression models (Supplementary Figure S5).

### Case study

To illustrate the use of models with censored covariates in differential discovery analysis, the FlowCAP IV dataset was reanalysed. A total of 766 flow cytometry PBMC samples linked to time to progression to AIDS from HIV+ of 383 patients (two per patient, one stimulated, one unstimulated) were available. For each sample, 13 marker intensities (IFN_*γ*_, TNF_*α*_, CD4, CD27, CD107-A, CD154, CD3, CCR7, IL2, CD8, CD57, CD45RO and V-Amine/CD14) together with channels FSC-A, FSC-H and SSC-A were measured. Of the 383 available survival times, 79 were observed, resulting in a censoring rate of 79% [25].

Preprocessing was performed according to the *FloReMi* pipeline: quality control, removal of margin events, doublet removal, compensation, logicle transformation and selection of alive T-cells [26].

*FlowSOM* was used for clustering with all marker intensities except FSC-C, FSC-H and SSC-A. The number of clusters was set to 400 and additionally, the metaclustering step was performed to obtain different subpopulation resolutions. The differential testing was then performed for a number of clusters of 20, 50, 100 and 400. The covariates were the survival time and the condition (stimulated or unstimulated) of the sample. Two random effects were modeled, one on a per sample level and the other on a per patient level. The three main methods (*rs*, *km*, *mrl*; number of imputations equal to 200) plus the complete case analysis were applied. An illustration of how the association between the survival time and the abundance for a cell population looks like is shown in Figure 6. At the top is a cluster with small adjusted p-value while the cluster in the bottom has a high adjusted p-value (as evaluated by *mrl*). No immediate association is visible, which could have various explanations, including high censoring rate, overdispersion, weak association.

**Figure 6.**
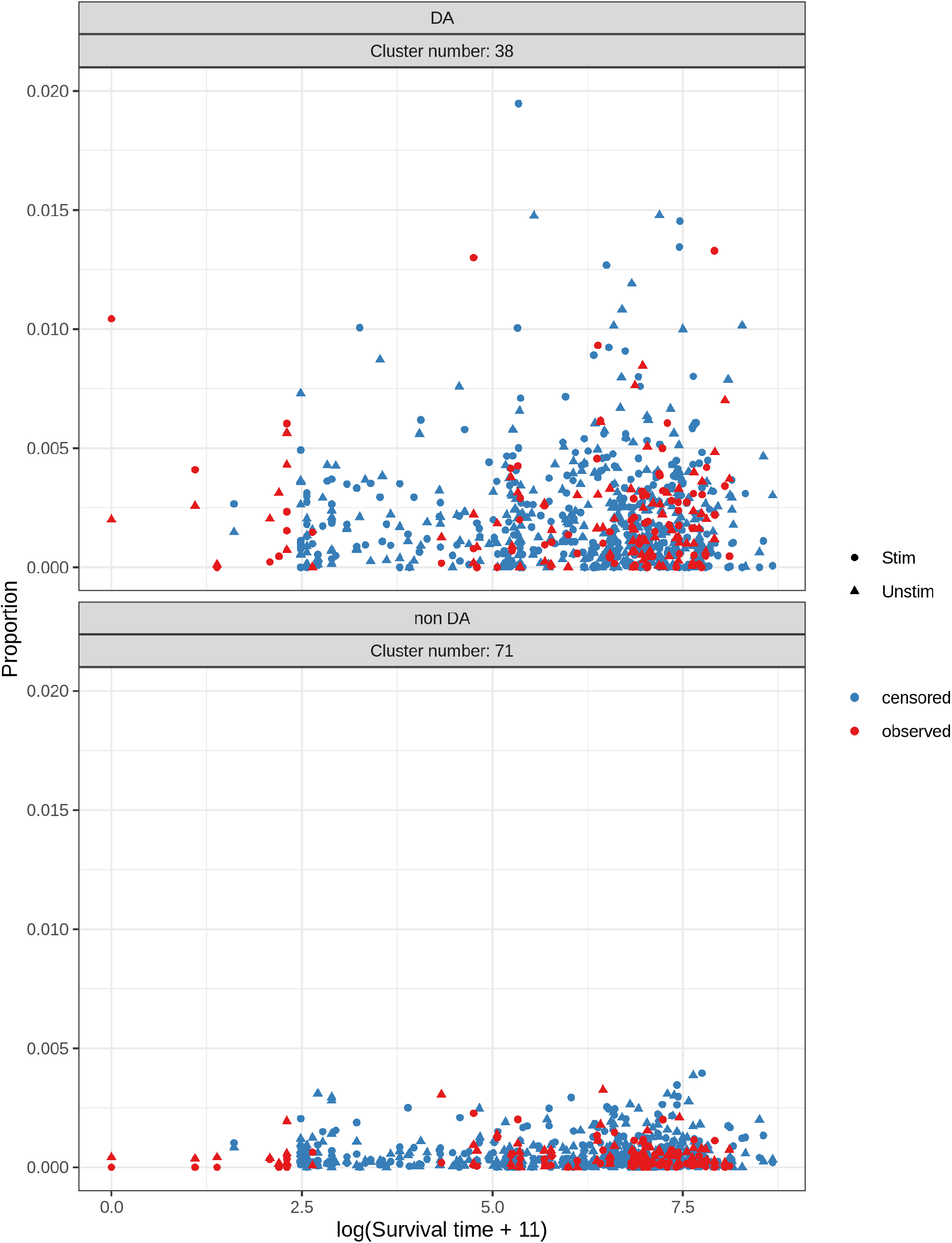
Association between cluster proportion and survival time for two clusters, one DA (top) and one non DA (bottom), evaluated with *mrl* for a total number of clusters of 100. The survival time is translated to get only positive values and then log transformed. Scaling of the axis in the upper plot removed 7 data points (0.9%).

Although no ground truth is established (i.e. which cell belongs to which cell population and which (if any) cell population is DA), a comparison to results from other methods (i.e. the original FlowCAP IV submissions) still gives insights into differential discovery performance. For the differential testing, the proportion of significant clusters for multiple cut offs differed substantially (Table 1). In general, the proportion of significant clusters is higher for a lower number of total clusters. While *rs* and *km* did not detect any DA clusters, *cc* found a large proportion of clusters to be significant and *mrl* has intermediate detection rates.

**Table 1.** Proportion of significant clusters for different total number of clusters (20, 50, 100, 200) for different significant levels (0.01, 0.05, 0.1) after multiple hypothesis correction in the case study. *cc*: complete case analysis, *km* Kaplan-Meier imputation, *mrl* mean residual life imputation (conditional multiple imputation), *rs* Risk set imputation.

| Number of cluster | cut off | mrl | rs | km | cc |
| --- | --- | --- | --- | --- | --- |
| 20 | 0.01 | 0.35 | 0 | 0 | 0.75 |
|  | 0.05 | 0.55 | 0 | 0 | 0.9 |
|  | 0.1 | 0.85 | 0 | 0.1 | 0.95 |
| 50 | 0.01 | 0.3 | 0 | 0 | 0.62 |
|  | 0.05 | 0.48 | 0 | 0 | 0.76 |
|  | 0.1 | 0.66 | 0 | 0 | 0.8 |
| 100 | 0.01 | 0.13 | 0 | 0 | 0.49 |
|  | 0.05 | 0.32 | 0 | 0 | 0.59 |
|  | 0.1 | 0.39 | 0 | 0 | 0.64 |
| 400 | 0.01 | 0 | 0 | 0 | 0.0575 |
|  | 0.05 | 0 | 0 | 0 | 0.07 |
|  | 0.1 | 0 | 0 | 0 | 0.108 |

Based on the proportion of detected clusters (Table 1), a level of 100 clusters was deemed to have a good balance between precision (cell population sizes) and sensitivity (proportion of detected clusters). A closer look at the (unadjusted) p-values of those clusters (at a level of 100 clusters) revealed similarities between the methods: 6 clusters were found in the 10 clusters with lowest (unadjusted) p-value for at least 3 methods. The adjusted p-values for *rs* and *km* are much higher than any reasonable significance level, however, *cc* and *mrl* have clusters that are differentially abundant. For *cc*, the proportion of significant clusters seems to be rather high (~50%), which is not unexpected given the poor error control observed in the simulations.

Comparing the marker expressions of those “top” 6 clusters (Supplementary Figure S6) with the discovered subpopulations in the FlowCAP IV challenge reveals some similarities. For example, Cluster 9 matches the described population of CD3+ CD4-CD14/VIVID+ CD57-cells [25]) and cluster 38 is similar to the CD4-D27-CD107a-CD154-CD45RO-population described in FloReMi [26].

## Discussion

In differential abundance analysis with a variable subject to censoring, existing methods make use of classical survival analysis methods, such as the Cox proportional hazard model. In particular, this would model the observed cell population abundances as predictors. The use of a reversed approach (cell population abundance as response), however, has the benefit to directly include confounders such as batch or age. The problem is that this reversed approach leads to a censored predictor, which renders standard differential abundance analysis inapplicable. A workaround to this issue is the use of multiple imputation, where the imputation step is specifically designed to handle censored values. Simulation studies indicate that in general, there is a gain by including the censored data instead of discarding samples (complete case analysis; *cc*) or treating censored values as missing (predictive mean matching; *pmm*).

More specifically, the basic simulations revealed consistent but slightly biased parameter estimation for the related methods *rs*, *km* and *kme*, and the simulations modeled from real data showed similar or increased performance in terms of sensitivity compared to *cc* but with better error control. Parameter estimation with *mrl* on the other hand was unbiased in the basic simulation, but the coverage rate was higher than expected, which typically leads to conservative performance. In the simulations modeled from real data, the conservative performance of *mrl* was apparent for low FDR, while the TPR was often (especially for higher censoring rates) higher than for other methods. In the case study (no ground truth), only *mrl* and *cc* were able to detect differentially abundant cell (sub) populations although especially for *cc*, the number of detected clusters was high, which could indicate many false positives. But since for *mrl* the FDR was in the simulations in general very low, this could mean that indeed many clusters are differentially abundant or alternatively, the real data is substantially different in structure compared to the simulations. For example, the simulations assumed a missing data mechanism that is missing-completely-at-random (MCAR), which might not be given in this case. Especially for *cc*, a missing data mechanism different from MCAR could be a problem since it is known to be biased under this condition. On the other hand, *mrl* (and *rs* and *km*) should be able to handle certain missing-at-random (MAR) cases [33], although this was not directly confirmed here.

The methods considered for direct inclusion of a censored covariate all rely on multiple imputation, which has the advantage of high interpretability since the underlying statistical models are classical GLMMs. A disadvantage are high computing costs caused by the need for repeated imputations (e.g. for high censoring rates, runtimes of 1 h instead of 1 min); runtimes can be nonetheless reduced through parallelization. The resolution at which to analyze is another issue, since a high number of clusters may reduce the statistical power imposed by multiple hypothesis correction, while associations with rare cell populations might be overlooked for a low total number of clusters. If a hierarchical structure of the cell populations is available (e.g. via metaclustering in *FlowSOM*), tree-based aggregated hypothesis testing methods (e.g. *treeclimbR* [35]) could increase differential discovery performance. Additional improvements of the differential discovery performance could be achieved by the use of a different analysis method such as *edgeR* or *voom*, which were shown to have increased performance compared to *GLMM* [18]. A further issue is of general nature: testing the association with a continuous (censored) covariate requires larger sample sizes compared to the testing with a binary covariate, although this nonetheless also depends on the dispersion and the strength of the association.

## Conclusion

Statistical modeling with a high proportion of censored data is always challenging, but even more so in DA settings with often overdispersed data and the need for multiple hypothesis testing correction. Nonetheless, we showed that including censored variables as a predictor in GLMMs results in high error control and decent sensitivity for a subset of the tested methods. Compared to classical survival analysis methods, such as the Cox proportional hazard model, higher flexibility in testing is provided, reflecting the need in typical experimental and clinical setups.

The tested methods were implemented in R and are submitted to Bioconductor. The source code is also available on GitHub https://github.com/retogerber/censcyt. Scripts for reproducing results and figures can be found on https://github.com/retogerber/censcyt_paper_scripts.

## Methods

### Censoring

The data mechanism for simulating censored data is based on the one described in Atem *et al.* [33]. The variable *X* to be censored is drawn from a Weibull distribution with scale *λ_x_* and shape *κ_x_* with the following parameterization:

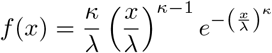

with the scale parameter *λ* > 0, the shape parameter *κ* > 0 and *x* ≥ 0. A second variable *C* that corresponds to the censoring time is also drawn from a Weibull distribution, but with different shape and scale parameters. The observed value *T* is then the minimum of *X* and *C*. In summary:

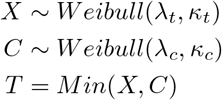

The parameters of the Weibull distributions are derived from the FlowCAP IV dataset [25]. More precisely *λ_x_* and *κ_x_* are obtained by fitting a Weibull distribution on the full dataset (taking into account censoring), while *λ_c_* is from fitting only on the censored samples. *κ_c_* is then calculated by first defining the desired censoring rate and then solving for *κ*_c_ (by calculating the probability 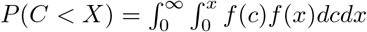, which can be seen as the expected censoring rate, for different values of *κ_c_*).

### Single cluster simulation

For the basic simulations with only a single cluster, the counts *Y_j_* (number of cells) with *j* ∈ 1*…N* was sampled from a generalized linear mixed model with a logit link function where the response (the number of cells) followed a binomial distribution (Eqn. 1) where *T_j_* follows a Weibull distribution with parameter as described above estimated from the FlowCAP IV dataset [25], the regression coefficients were set to *b*_0_ = −2, *b*_1_ = −0.0001 and *b*_2_ = 1, *Z_j_* ∈ {0, 1} is a binary covariate with balanced groups, *R_j_* is an observation level random effect to model overdispersion distributed according to a standard normal distribution (*σ*^2^ = 1) and *n_j_* is the sample size distributed according to a uniform distribution with a minimum limit of 10’000 and a maximum limit of 100’000.

### Multiple cluster simulation

The matrix of counts *Y* ∈ *R^K×N^* for *K* clusters (cell populations) and *N* samples follows a Dirichlet-Multinomial (DM) distribution (Eqn. 2) for *j* ∈ {1*…N*} where *n_j_* is the total number of cells in sample *j* and 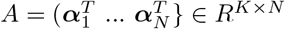 with *A_ij_* > 0 for *i* ∈ {1*…N*} are the concentration parameters dependent on covariates *T_j_* and *Z_j_*. Additionally *Y_j_* = (*Y*_1*j*_…*Y_kj_*), *T_j_ ~ Weibull*(*λ, κ*) and *Z_j_* ∈ {0, 1} is a binary variable with balanced groups. The proportions of cells in cluster *i* in sample *j* is simply

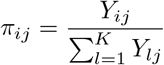

An association for cluster *i* is then assumed to be the following:

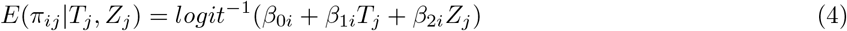

with an intercept *β*_0*i*_, a slope *β*_1*i*_ for *T_j_* and a slope *β*_2*i*_ for *Z_j_*. The *β*’s are therefore fixed for a cluster but are different between clusters. The covariates *T_j_* and *Z_j_* are specific for a sample but not a cluster. The proportions *π_j_* for sample *j* follow a Dirichlet distribution, meaning the *π_ij_* themselves follow a Beta distribution with mean

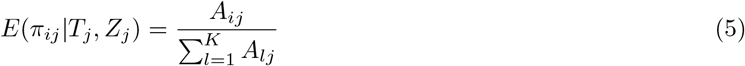

This allows to combine Eqn. 4 and Eqn. 5 leading to Eqn. 6 (which is the same as Eqn. 3):

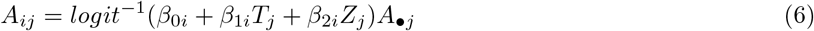

with the sum of the concentration parameters for a sample 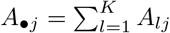. This means that *A_ij_*|*T_j_, Z_j_* is dependent on six values: the intercept *β*_0*i*_, the first slope *β*_1*i*_, the continuous covariate *T_j_*, the second slope *β*_2*i*_, the binary covariate *Z_j_* and *A_•j_*. Since the sum *A_•j_* depends on all *A_ij_* for a given sample *j*, this means that in order to keep this sum equal across samples, for every *A_ij_* that is increased with increasing *T_j_* there has to be an *A_ij_* that decreases the same amount. Because of the non-linearity of the *logit* function this could lead to a very weak association of the second *A_ij_* (which would not strictly follow the logit relationship). The strategy is therefore to allow small discrepancies of the sum *A_•j_* in order to get the specified associations. To decrease the variation of the sum *A_•j_* two clusters of similar proportion are chosen. To obtain *β*_0_ and *β*_1_, the desired minimum / maximum mean proportion *π_ij_* for *max*(*T_j_*) is determined and then *Z_j_* is set to zero to solve for *β*_0_ and *β*_1_. This will result in a sum *A_•j_* that is exactly the same at *T_j_* = 0 and *T_j_* = *max*(*T_j_*). All sums *A_•j_* in between will slightly deviate but this deviation is too small to detect under the simulation conditions considered here. To obtain *β*_2*i*_, a difference of the mean abundance at *T_j_* = 0 is specified, which then allows to calculate *β*_2*i*_. In short, the *β*’s are calculated by specifying border constraints, consisting of maximum differences in the mean abundance dependent on the covariates. Because it was observed that the spread of the simulated data was higher than in the real dataset, the concentration parameters were multiplied by a factor of five (keeping the expected counts per cluster the same) to reduce the variance of counts.

### Multiple imputation

The goal of multiple imputation is not to replace the missing or censored values by estimates but rather to find a parameter estimate of the statistical model being tested that is *unbiased and confidence valid* [34].

Multiple imputation consists of three main steps [34]: Imputation, Analysis, Pooling. In the first step, multiple complete datasets are generated by replacing the incomplete values with a random draw from a set of possible true values. This can, for example, be the assumed or empirical distribution of the incomplete value. In the second step, each completed dataset is individually analysed, e.g. by fitting a regression model. In the third step, the results from the second step are combined using Rubin’s rules [36] that consider the additional variances in the analysis. A slight variation is the use of *Resampling* in the first step. Before imputation, a bootstrap sample is drawn, which is then the new incomplete dataset where the missing values get replaced. One of the advantages of this approach: the incomplete value can be replaced by a deterministic quantity of the data (e.g. the mean), which would not work in classical multiple imputation (each imputed dataset would be the same). A drawback is that *Resampling* techniques are based on large-sample theory and might not work properly for small samples [29].

### DA

The presented DA methods are based on the GLMM approach in *diffcyt* which consists of fitting a generalized linear mixed model with a logit link function for each cell population, testing and multiple hypothesis testing correction. When a censored covariate is present, multiple imputation is used to handle the additional uncertainties of the parameters caused by incomplete data. The imputation methods are described in the following.

In complete case analysis (*cc*, also known as listwise deletion [34]), only the observed values (***T** _j_*|***T** _j_ < **C**_j_* for *j* ∈ {1*…N*}) are used by discarding all incomplete samples.

The risk set imputation (*rs*) [31] first constructs the risk set *R*(***T** _l_*) = {***T** _j_*|***T** _j_ > **T** _l_*} for *j* ∈ {1*…N*} for all ***T** _l_ **T** _l_ < **X**_l_* with *l* ∈ {1*…N*} and second, randomly draws one of those as the imputed value. If censoring depends on a covariate, the risk set is calculated as described in (Hsu *et al.*) [37], incorporating the idea of predictive mean matching.

The Kaplan-Meier imputation (*km*) [31] is similar to risk set imputation. It first constructs the risk set *R*(***T** _l_*) for all ***T*** _*l*_|***T*** _*l*_ < ***X***_*l*_ and then estimates the survival function with the Kaplan-Meier estimator for each of those sets. A random event time according to the survival curve is drawn and replaces the censored value.

Conditional multiple imputation [33] (labeled here as mean residual life imputation (*mrl*)) is based on the mean residual life, which is the expected remaining survival time until an event happens

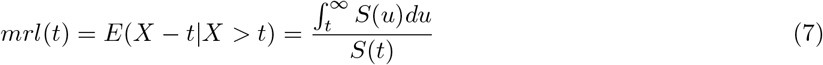

with the random variable *X* representing the true (unobserved) survival time and *S*(*t*) the survival function. It can be used to get an estimate of how long it will take until an event happens given that the event did not happen yet. Conditional single imputation [33] (Conditional multiple imputation with only one imputation) imputes censored values by adding the corresponding mean residual life. First a survival curve *S*(***T***) (using the Kaplan-Meier estimator) is fitted and then the mean residual life is added to the censored value [33]. If censoring depends on a covariate, *S*(***T***) can be fitted using the Cox-proportional hazards model [27]. Mean residual life imputation (Conditional multiple imputation) can not be used in the normal multiple imputation set up since all imputed datasets would be the same. Instead *Resampling* is applied to first generate incomplete datasets before imputation.

The estimation of *S*(***T***) is done without any distributional assumptions resulting in a high data dependency. If the sample size is small and/or many values are censored the estimation can be drastically different from the true (unobserved) survival function. Especially towards the tails, as data gets even sparser, estimation is difficult. If the highest measured value is censored, *S*(***T***) does not reach its theoretical minimum (zero). The usual way to deal with this problem is to treat the maximum value as if it was observed. Another possibility is to make a distributional assumption for the tail of the survival function. This was explored for the method Kaplan-Meier imputation by assuming an exponential tail, which is referred to here as *kme* (based on [38]).

Unfortunately, there is no clear rule as to how many imputations are needed [34]. In general, this depends on (among other things) the censoring rate; higher censoring requires more imputations. Two methods to estimate the number of imputations are based on a linear rule [39] and a quadratic rule [40]. Only minor changes in the results after around 50 imputations could be seen in our case leading to the use of 50 imputations as the default.

### Case study

Following are some clarifications of the description in the main text. The raw flow cytometry data is available under http://flowrepository.org/id/FR-FCM-ZZ99. The data set consists of 766 PBMC samples linked to time to progression to AIDS from HIV+ of 383 patients. For each patient, two samples (measured at the same time) are available: one untreated and one treated with HIV-Gag proteins.

Preprocessing was performed according to the *FloReMi* pipeline [26]: First, quality control by removing cells within a certain time sampling interval where the median FSC-A value differed dramatically from tolerable limits. Then, removal of margin events by removing cells that have a minimum and maximum value for some channel. Next, the selection of single cells by removing cells whose FSC-A to FSC-H ratio was larger than the median ratio plus two times the standard deviation of the ratios. Next, compensation with the given spillover matrices (from the .fcs files) was applied, data was logicle transformed and alive T-cells were gated (using *flowDensity*) using channels V-Amine/CD14 and CD3 and selection of V-Amine/CD14-CD3+ population.

In the differential testing a transformed survival time, according to *s_trans_* = *log_e_*(*s* + 11) (the +11 is to obtain only positive values since the lowest survival time is −10), was used.

## Acknowledgements

The authors thank Lukas M. Weber for help with the implementation and authorization for re-usage of code, and Stephanie Leemann for help designing Figure 3.

## Abbreviations

scRNA-seq: single-cell RNA sequencing
DA: differential abundance
DS: differential state
GLMM: generalized linear mixed model
RB: raw bias
CR: coverage rate
CI: confidence interval
RMSE: root mean squared error
km: Kaplan-Meier imputation
kme: Kaplan-Meier imputation with an exponential suvival function tail
rs: risk set imputation
mrl: mean residual life imputation
pmm: predictive mean matching
cc: complete case analysis
DM: dirichlet-multinomial distribution
TPR: true positive rate
FDR: false discovery rate

## Ethics approval and consent to participate

Not applicable.

## Consent for publication

Not applicable.

## Authors’ contributions

RG and MDR developed methods, designed analyses, and wrote the manuscript. RG implemented methods and performed analyses. All authors read and approved the final manuscript.

## Additional Files

**Additional file 1 — Supplementary figures**

Additional Figures, as mentioned in the main text.

