## Supplementary Figures for "censcyt: censored covariates in differential abundance analysis in cytometry"

November 2020

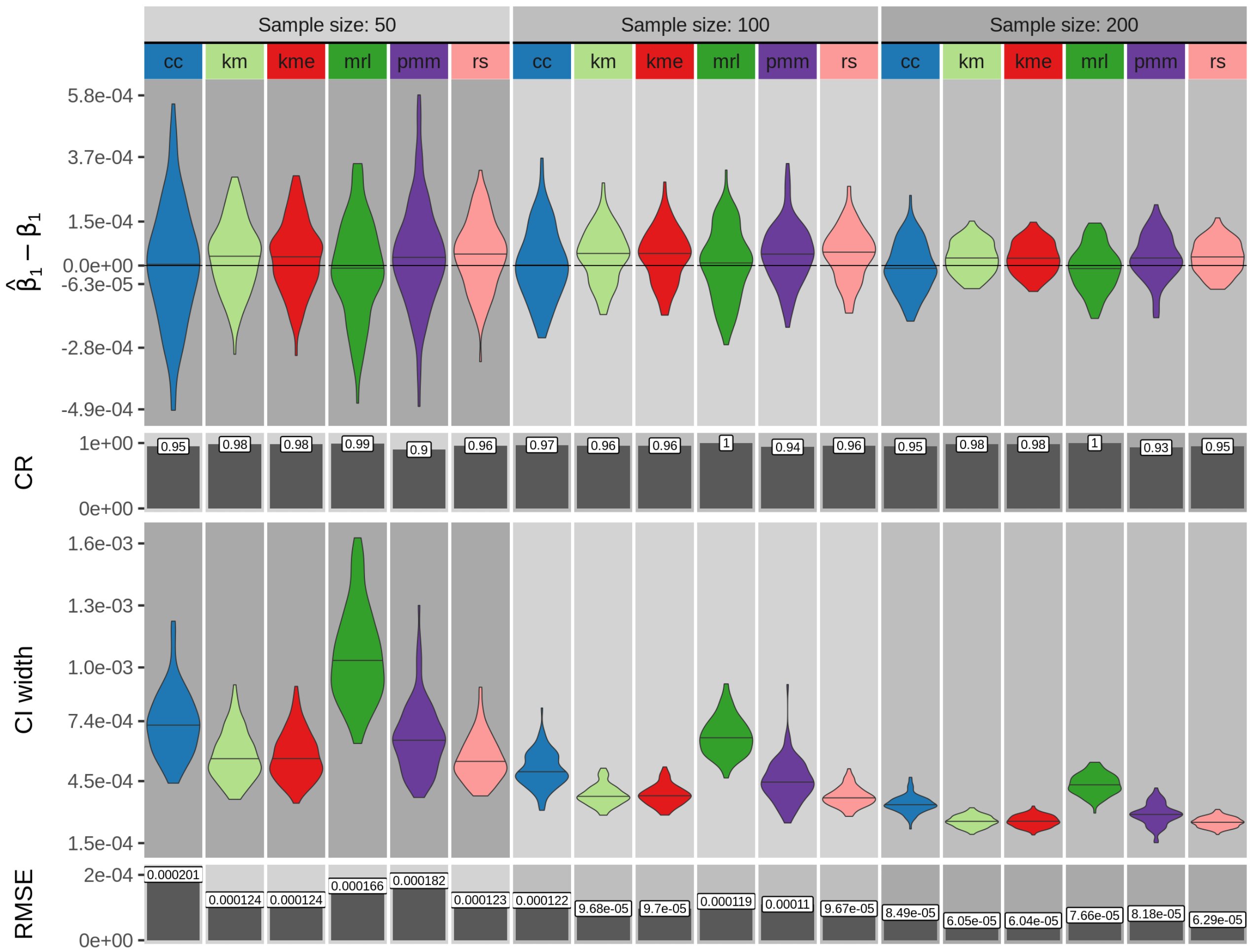


Figure S1: Single cluster simulation results for a censoring rate of 50% for sample sizes of 50, 100 and 200. Shown are four measures calculated from 100 simulation repetitions: difference of the estimated regression coefficient ($\hat{\beta}_1$) and its true value ($\beta_1$) , coverage rate (CR), confidence interval (CI) width and root mean squared error (RMSE). *cc*: complete case analysis, *km*: Kaplan-Meier imputation, *kme*: Kaplan-Meier imputation with an exponential tail, *mrl*: mean residual life imputation (conditional multiple imputation), *pmm*: predictive mean matching (treating censored values as missing), *rs*: risk set imputation. Other parameter values are: true regression coefficient $\beta_1 = -1e-4$, number of multiple imputations = 50 and the variance of the random effect = 1.

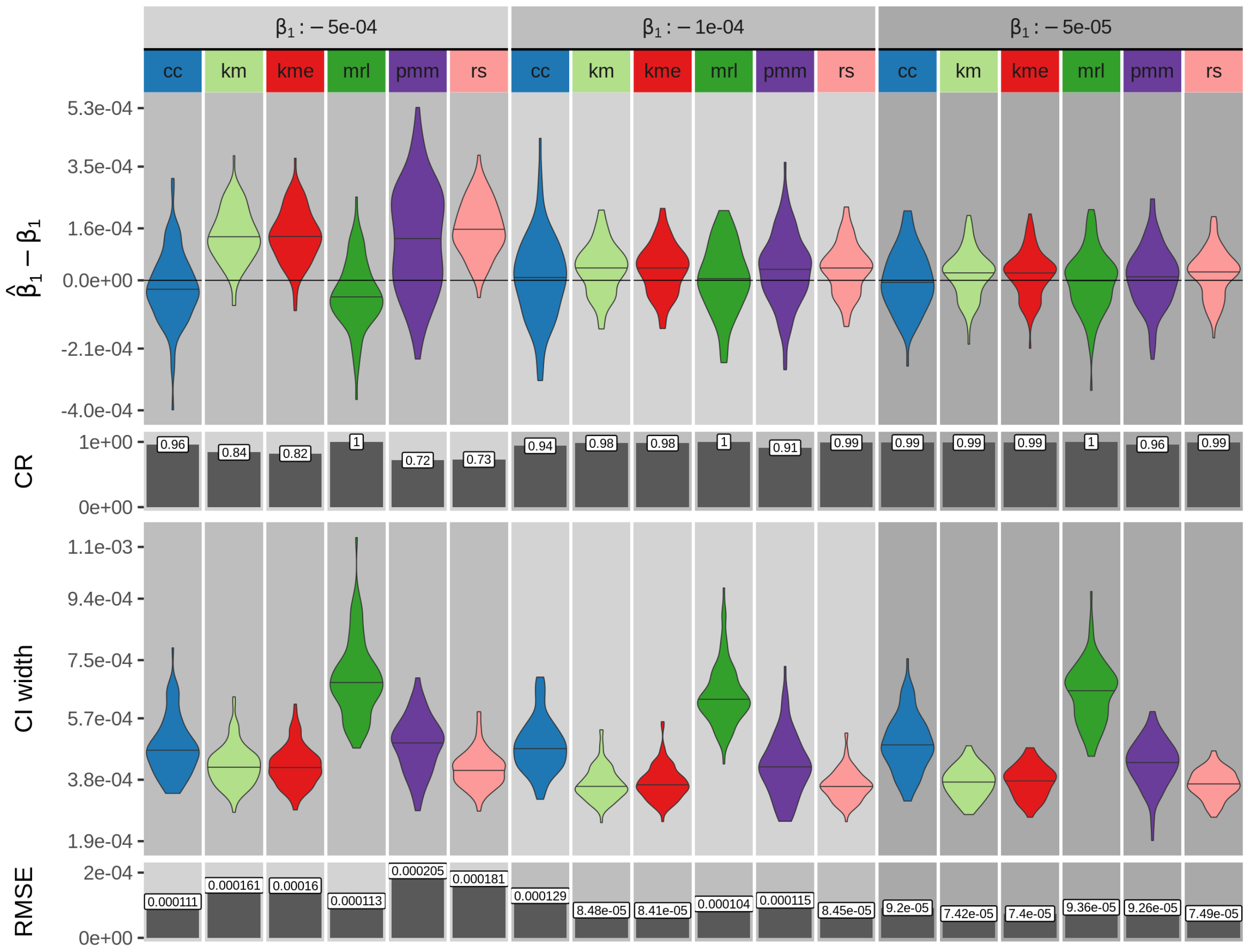


Figure S2: Single cluster simulation results for a sample size of 100 and a censoring rate of 50% for regression coefficients $\beta_1$ of $-5e-5$,$-1e-4$ and $-5e-4$. Shown are four measures calculated from 100 simulation repetitions: difference of the estimated regression coefficient ($\hat{\beta}_1$) and its true value ($\beta_1$) , coverage rate (CR), confidence interval (CI) width and root mean squared error (RMSE). *cc*: complete case analysis, *km*: Kaplan-Meier imputation, *kme*: Kaplan-Meier imputation with an exponential tail, *mrl*: mean residual life imputation (conditional multiple imputation), *pmm*: predictive mean matching (treating censored values as missing), *rs*: risk set imputation. Other parameter values are: number of multiple imputations = 50 and the variance of the random effect = 1.

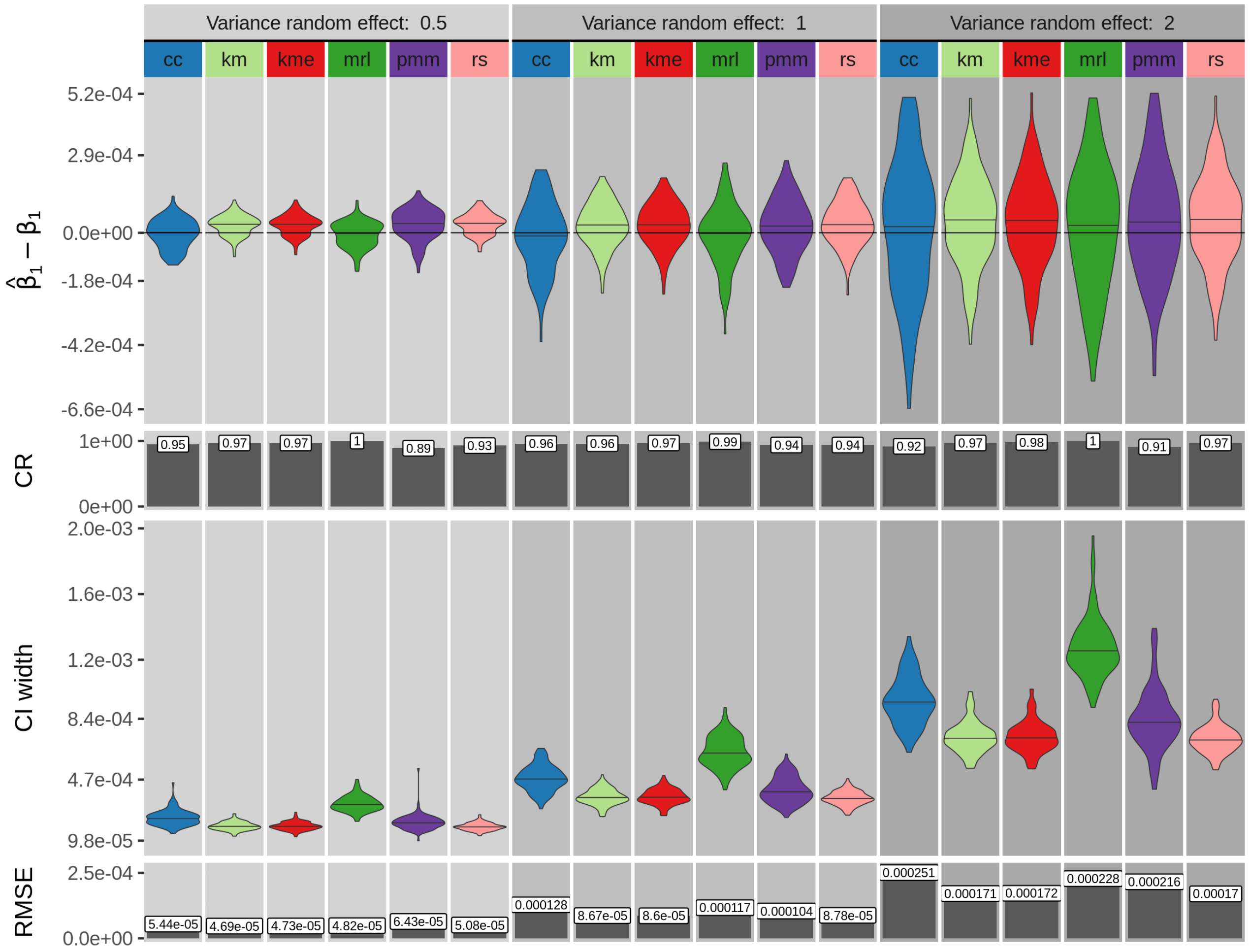


Figure S3: Single cluster simulation results for a sample size of 100 and a censoring rate of 50% for variances of the random effect of 0.5, 1 and 2. Shown are four measures calculated from 100 simulation repetitions: difference of the estimated regression coefficient ($\hat{\beta}_1$) and its true value ($\beta_1$) , coverage rate (CR), confidence interval (CI) width and root mean squared error (RMSE). *cc*: complete case analysis, *km*: Kaplan-Meier imputation, *kme*: Kaplan-Meier imputation with an exponential tail, *mrl*: mean residual life imputation (conditional multiple imputation), *pmm*: predictive mean matching (treating censored values as missing), *rs*: risk set imputation. Other parameter values are: true regression coefficient $\beta_1 = -1e-4$ and number of multiple imputations = 50.

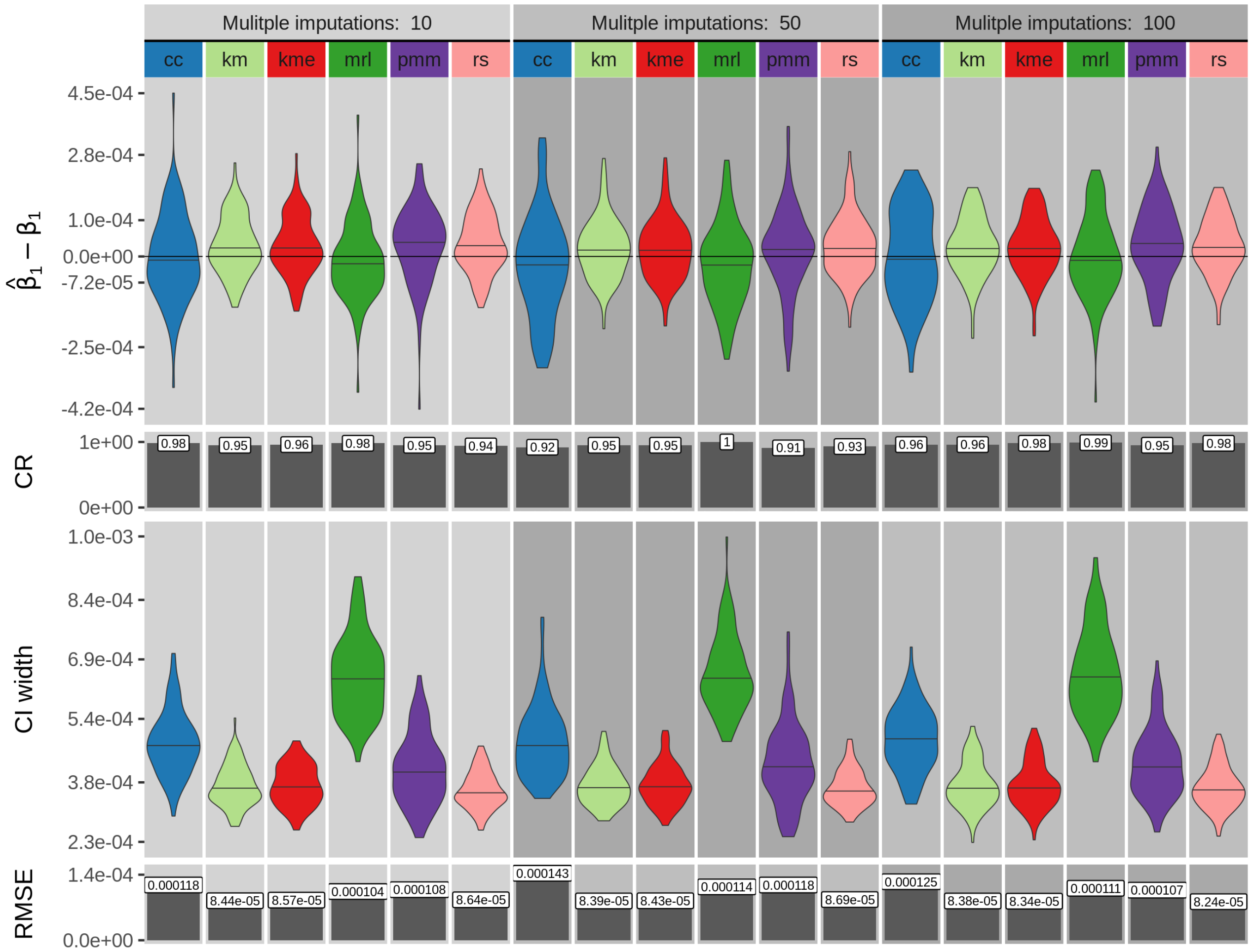


Figure S4: Single cluster simulation results for a sample size of 100 and a censoring rate of 50% for number of imputations of 10, 50 and 100. Shown are four measures calculated from 100 simulation repetitions: difference of the estimated regression coefficient ($\hat{\beta}_1$) and its true value ($\beta_1$) , coverage rate (CR), confidence interval (CI) width and root mean squared error (RMSE). *cc*: complete case analysis, *km*: Kaplan-Meier imputation, *kme*: Kaplan-Meier imputation with an exponential tail, *mrl*: mean residual life imputation (conditional multiple imputation), *pmm*: predictive mean matching (treating censored values as missing), *rs*: risk set imputation. Other parameter values are: true regression coefficient $\beta_1 = -1e-4$ and the variance of the random effect $= 1$.

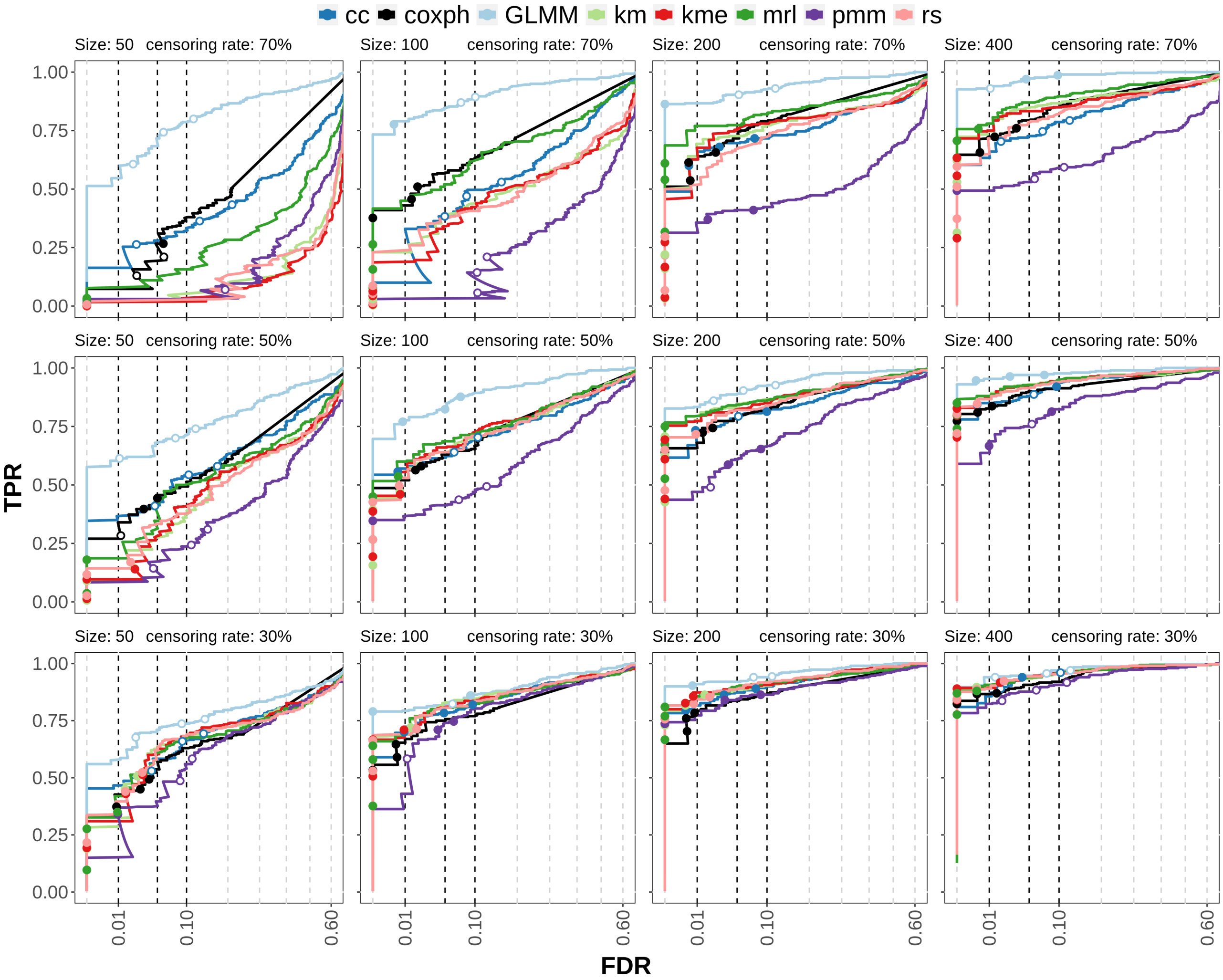


Figure S5: Multiple cluster simulation results testing with only a single (censored) covariate. TPR-FDR curves for censoring rates of 30%, 50% and 70% (rows) and samples sizes of 50,100,200 (columns). The x-axis is square root transformed. *cc*: complete case analysis, *km*: Kaplan-Meier imputation, *kme*: Kaplan-Meier imputation with an exponential tail, *mrl*: mean residual life imputation (conditional multiple imputation), *pmm*: predictive mean matching (treating censored values as missing), *rs*: risk set imputation, *coxph*: Cox proportional hazards model. *GLMM* uses the (unobserved) ground truth of the survival time and can be considered to be the maximum possible performance of the other methods.

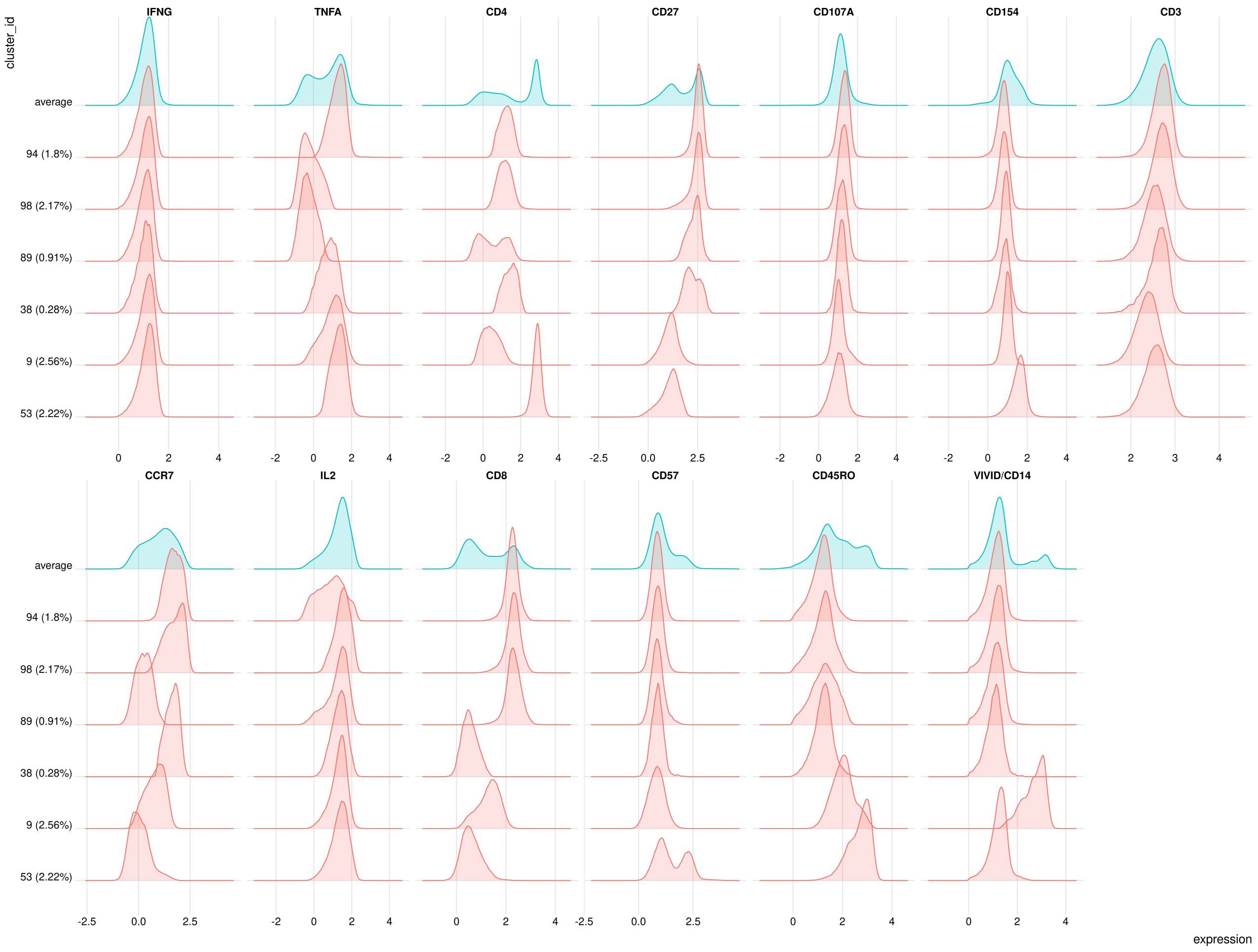


Figure S6: Expressions of top subpopulations of the clustering at a resolution of 100 clusters in the case study. The top row ("average") is the average over all 100 clusters.
